# Tracing infant sleep neurophysiology longitudinally from 3 to 6 months: EEG insights into brain development

**DOI:** 10.1101/2025.06.25.661476

**Authors:** Matthieu Beaugrand, Valeria Jaramillo, Christophe Mühlematter, Sarah F. Schoch, Vivien Reicher, Andjela Markovic, Salome Kurth

**Affiliations:** University of Fribourg, Department of Psychology, Fribourg, Switzerland; University of Surrey, Surrey Sleep Research Centre, Guildford, United Kingdom; University Hospital Zurich, Department of Pulmonology, Zurich, Switzerland; Donders Institute for Brain, Cognition and Behaviour, Radboud University Medical Centre, Nijmegen, NL; Department of Ethology, ELTE Eötvös Loránd University, Budapest, Hungary; Research Centre for Natural Sciences Institute of Cognitive Neuroscience and Psychology, Clinical and Developmental Neuropsychology Research Group, Budapest, Hungary; Semmelweis University, Department of Psychiatry and Psychotherapy, Budapest, Hungary; Translational Research Center, University Hospital of Psychiatry and Psychotherapy, Bern

**Keywords:** developmental trajectory, synaptic plasticity, ontogeny, signal processing, functional connectivity

## Abstract

Sleep is critical for brain plasticity during early development, yet the individual maturation of sleep neurophysiology in infancy remains poorly characterized. In particular, slow wave activity (SWA) has emerged as a key marker of both cortical maturation and experience-dependent plasticity. Understanding the regional dynamics of sleep neurophysiology early in life could yield critical insights into neurodevelopmental health. We conducted a longitudinal high-density EEG study in 11 healthy infants (3-6 months) assessing non-rapid eye movement (NREM) sleep. We analyzed the maturation of SWA (0.75-4.25 Hz), theta power (4.5-7.5 Hz), and sigma power (9.75-14.75 Hz) across scalp regions and examined their association with behavioral development. From 3 to 6 months, SWA increased maximally in occipital regions, while theta power exhibited a global increase. Sigma power, initially concentrated centrally, dispersed towards frontal regions. Greater power increases over frontal regions correlated with higher motor (theta) and personal-social skill scores (sigma) at 6 months. These findings establish a framework for typical infant sleep EEG maturation, highlighting frequency-specific and regionally distinct developmental patterns. This study provides the first longitudinal evidence that early changes in sleep EEG topography reflect individual developmental trajectories, supporting its utility as a non-invasive and yet precise biomarker for early identification of atypical neurodevelopment at preverbal ages.

## 2. Introduction

At birth, the human brain size has reached only 27% of its adult size (Robson & Wood, 2008), and the postnatal refinement of cortical microstructure progresses rapidly through processes such as synaptogenesis and myelination. Cortical development follows a temporally staggered trajectory, with sensorimotor regions maturing earlier than association cortices, as shown by longitudinal imaging studies of metabolic activity (Chugani et al., 1987), gray matter growth (Gogtay et al., 2004), and cortical thickness (Shaw et al., 2008). Infancy thus represents a critical period for brain maturation, during which sleep is believed to play a central role in neuronal development. Sleep disruptions during this period can have lasting consequences and disturbed sleep neurophysiology accompanies a multitude of pathophysiologies in children, adolescents, and adults (Markovic et al., 2021; Mouthon et al., 2016; Tesler et al., 2016), for a review, see (Tarokh et al., 2016). The sleep electroencephalogram (EEG) provides a non-invasive window into these neurofunctional processes, revealing region-specific changes in neural activity. Sleep architecture and functional brain networks identified through the sleep EEG carry critical insights into brain health throughout the lifespan, with essential physiological indicators of neuronal wellbeing already in newborns (Tokariev et al., 2019).

Three key sleep EEG features – slow wave activity (SWA; 0.75-4.25 Hz), theta power (4.5-7.5 Hz), and sigma power (9.75-14.75 Hz) – serve as markers of cortical maturation. SWA, a hallmark of non-rapid eye movement (NREM) sleep, signals synaptic homeostasis (Tononi & Cirelli, 2006) and local neuroplasticity, with learning-related increases shown in both children and adults (Fattinger et al., 2017; Huber et al., 2004; Wilhelm et al., 2014). SWA undergoes marked transformations in amplitude, morphology, and topographical distribution throughout childhood and adolescence (Campbell et al., 2011; Jaramillo et al., 2020; Kurth, Jenni, et al., 2010; Tarokh & Carskadon, 2010), with its dominant expression shifting from posterior to anterior scalp regions (Kurth, Ringli, et al., 2010a). These changes mirror patterns of gray and white matter maturation (Buchmann et al., 2011; LeBourgeois et al., 2019) and reflect underlying synaptic remodeling and homeostasis (Huttenlocher & Dabholkar, 1997; Timofeev et al., 2020)(Tononi & Cirelli, 2006). SWA topography is associated with cognitive outcomes (Buchmann et al., 2011; Ednick et al., 2009; Reynaud et al., 2018) and its regional shifts predict brain function, preceding motor skill emergence by several years (Kurth et al., 2012).

Deviations from typical SWA distribution are evident in neurodevelopmental and neurological conditions such as ADHD, epilepsy and depression (Eriksson et al., 2023; Furrer et al., 2019; Tesler et al., 2016), further highlighting dual roles of SWA in marking of cortical maturity and functional plasticity. Yet, SWA maturation during infancy – when plasticity is highest – remains poorly understood.

Theta power (4.5-7.5 Hz) in sleep EEG provides key insights into neurophysiological maturation, complementing SWA. It undergoes substantial developmental changes from infancy to adolescence (Novelli et al., 2016; Otero et al., 2011; Page et al., 2018) and serves as a transitional marker of sleep homeostasis (Campbell et al., 2011; Jenni et al., 2004). Atypical theta patterns are linked to neurodevelopmental disorders (Gorgoni et al., 2020), with increased theta power in ADHD (Castelnovo et al., 2022), reduced fronto-central connectivity in Asperger’s Syndrome (Lázár et al., 2010), and widespread hyperconnectivity in childhood-onset schizophrenia (Markovic et al., 2021).

Sigma power (9.75-14.75 Hz), linked to sleep spindles in NREM sleep, exhibits marked developmental changes. Sleep spindles are bursts of activity in the sigma frequency range generated through thalamocortical loops (Sanchez-Vives & McCormick, 2000; Steriade, 2006). In early childhood, spindles increase in duration and amplitude but decrease in frequency (McClain et al., 2016), while adolescence brings higher spindle density and frequency with lower amplitude and duration (Goldstone, Willoughby, de Zambotti, et al., 2019; Hahn et al., 2019). During development, sigma power extends from a central focus into both frontal and parietal regions (Goldstone, Willoughby, de Zambotti, et al., 2019; Kurth, Ringli, et al., 2010a; Tarokh & Carskadon, 2010). Sleep spindles play a crucial role in memory consolidation (De Gennaro & Ferrara, 2003) and cognitive development (Goldstone, Willoughby, De Zambotti, et al., 2019; Hoedlmoser et al., 2014), with early spindle activity predicting psychomotor skills (Satomaa et al., 2020) and behavioral outcomes (Jaramillo et al., 2023). Clinically, sigma power is reduced in childhood-onset schizophrenia and correlates with symptom severity (Markovic et al., 2020), while lower levels are linked to depressive symptoms in adolescents (Hamann et al., 2019), underscoring its indicative role in broader neuropsychological conditions.

Altered sleep EEG oscillations in neurodevelopmental disorders can further impair cognitive maturation (Gorgoni et al., 2020), highlighting EEG oscillatory activity as a potential treatment target (Bölsterli et al., 2011). To develop effective interventions, it is essential to establish normative sleep EEG topography across infancy and childhood, as patterns evolve dynamically with age and region (Dereymaeker et al., 2017; Kurth, Ringli, et al., 2010b). Sleep EEG topography mapping provides area-specific insights into neurodevelopment (Page et al., 2018; Satomaa et al., 2020), and tracking early EEG trajectories across frequency bands and brain regions, could yield normative trajectories and non-invasive biomarkers (Mason et al., 2021). Combining multiple parameters – SWA, theta, and sigma power – may enhance diagnostic precision beyond single-metric analyses (Baglioni et al., 2010).

Longitudinal within-subject sleep EEG studies in early life provide a unique opportunity to track brain activity maturation, map neural plasticity processes, and assess their influence on later behavioral outcomes. This study aimed to characterize the typical topographical maturation of infant sleep EEG from 3 across 6 months. We hypothesized that SWA would show the greatest increase in occipital regions, theta power would develop prominently over both occipital and central regions, and sigma power would be concentrated centrally. By examining regionally and frequency-specific EEG changes over time, we aimed to further capture early trajectories of brain maturation and their predictive value for behavioral outcomes during this highly sensitive developmental window.

## 3. Methods

### 3.1. Participants and study design

This study included 11 healthy infants (3 girls) who met strict inclusion criteria. All were born at term via vaginal delivery with a birth weight exceeding 2500g. Families were required to be native or highly proficient French speakers. Exclusion criteria were central nervous system disorders, brain injuries, chronic illnesses, or a family history of sleep or mental health disorders. The study was approved by the cantonal ethics committee and conducted in accordance with the Declaration of Helsinki. Parents provided written informed consent after receiving a detailed explanation of the study procedures.

Before and at enrollment, all infants were required to be in good health and free from medications, including antibiotics, that could affect the sleep-wake cycle. High-density sleep EEG recordings were conducted at home at 3 months (n=9) and 6 months (n=11). While most infants participated at both time points, two were only assessed at 6 months. Developmental outcomes were measured using the parent-rated Ages and Stages Questionnaire (Squires et al., 1995), completed within 7 days before or after each EEG assessment.

### 3.2 Sleep EEG

High-density EEG recordings were conducted during the infants’ regular bedtime to maintain a natural sleep environment. A 124-electrode sponge net (Electrical Geodesics Sensor Net, Electrical Geodesics Inc., EGI, Eugene, OR) was prepared by soaking it in an electrolyte solution (1 L warm tap water, 10 mL potassium chloride, 1 mL baby shampoo) for 3-5 minutes. The net was carefully adjusted to each infant’s head, aligned with the vertex and mastoids. EEG data were recorded using Brain Vision Recorder (v 1.25.0201, Brain Products GmbH, Gilching, Germany) at 500 Hz sampling frequency, referenced to the vertex (Cz), with impedances kept below 50 kΩ. Each session lasted up to two hours.

Raw EEG data were preprocessed according to in-lab standards (Kurth, Ringli, et al., 2010a; Schoch et al., 2022). Signals were filtered with a 0.5-50 Hz bandpass filter and down-sampled to 128 Hz. Sleep stages were scored in 20-second epochs by two independent scorers following the American Academy of Sleep Medicine Manual (AASM) with pediatric adjustments (Iber, 2007), with discrepancies resolved by consensus. Artifacts were identified semi-automatically by analyzing power levels across frequency ranges relative to a predefined threshold (Huber et al., 2000). Channels with poor signal quality (e.g., muscle artifacts on outermost electrodes near the neck/face) were excluded. Each channel’s signal was re-referenced to the average of all channels (i.e., average reference). EEG power was quantified across SWA (0.75-4.25 Hz), theta (4.5-7.5 Hz), and sigma (9.75-14.75 Hz) bands, averaging across the remaining channels. Analyses focused on the first 60 minutes of artifact-free NREM sleep, aiming to include the maximal available duration within this window.

### 3.3 Behavioral development

Infant development was assessed using an age- and language-adapted version of the Ages and Stages Questionnaire (ASQ), completed by primary caregivers (Squires et al., 1995). This widely used tool evaluates five developmental domains: Communication, Gross motor, Fine motor, Problem solving, and Personal social, along with a composite Collective score (Gollenberg et al., 2010).

For this study, we focused on the Gross motor and Personal social domains, which are particularly relevant to early infancy. Gross motor skills assess muscle use in activities such as rolling, sitting, and walking, with delays among the most common early developmental challenges (Valla et al., 2015). Prior research has shown that sleep spindle density at 6 months predicts Gross motor outcomes at 12 and 24 months (Jaramillo et al., 2023).

The Personal social subdomain evaluates infants’ interactions and emerging self-care behaviors. Our previous research with over 150 infants found a correlation between sleep patterns and personal social development (Schoch et al., 2022), supporting evidence that sleep plays a key role in socio-emotional growth (Kaley et al., 2012; Mindell et al., 2017; Williams et al., 2016).

### 3.4 Statistical Analysis

Paired t-tests were performed at 0.25-Hz intervals from 0.25 to 19 Hz to assess sleep EEG power maturation across the frequency spectrum. These tests analyzed power values averaged across subjects in key electrodes of standard EEG set ups: F3, C3, P3, O1 (left hemisphere), and F4, C4, P4, O2 (right hemisphere), chosen for their established relevance in sleep EEG research along the anterior-posterior axis.

To examine topographical changes in SWA, theta power, and sigma power between 3 and 6 months, paired t-tests were conducted at each electrode for infants with complete data at both time points.

Within-subject development was quantified by calculating power differences for each electrode, subtracting 3-month values from 6-month values. To explore associations between EEG maturation and behavioral development, Spearman correlations were computed between maturational changes in power and Collective, Gross motor, and Personal social scores at 6 months.

Permutation tests were applied to all correlations and t-tests to ensure robust statistical inference. For each test, an empirical p-value was computed by comparing the absolute value of the observed test statistic to a null distribution of absolute test statistics obtained through 1000 permutations of the original data. Specifically, the p-value reflects the proportion of permuted values whose magnitude exceeded that of the observed statistic. A result was considered statistically significant if this p-value was below 0.05. To maintain error control, the permutation structure (i.e., the shuffling order) remained consistent across electrodes and frequencies throughout 1000 iterations.

## 3. Results

The duration of artifact-free NREM sleep included in the analyses ranged from 28 to 60 minutes at the 3-month assessment (mean ± SD: 50 ± 12 minutes) and from 33 to 56 minutes at the 6-month assessment (mean ± SD: 49 ± 8 minutes). We first investigated the maturational dynamics of each hemisphere by analyzing power spectra at selected electrodes (Figure 1). In the left hemisphere, the frontal electrode (F3) showed a significant increase in power from 3 to 6 months, particularly in the 1-2 Hz and 4-10 Hz ranges. In contrast, the central electrode (C3) displayed no significant change. The parietal electrode (P3) exhibited a significant power increase between 2-6 Hz. In contrast, the occipital electrode (O1) revealed the most pronounced changes, with a widespread increase in power across nearly all frequencies above 1 Hz (Figure 1A).

**Figure 1.**
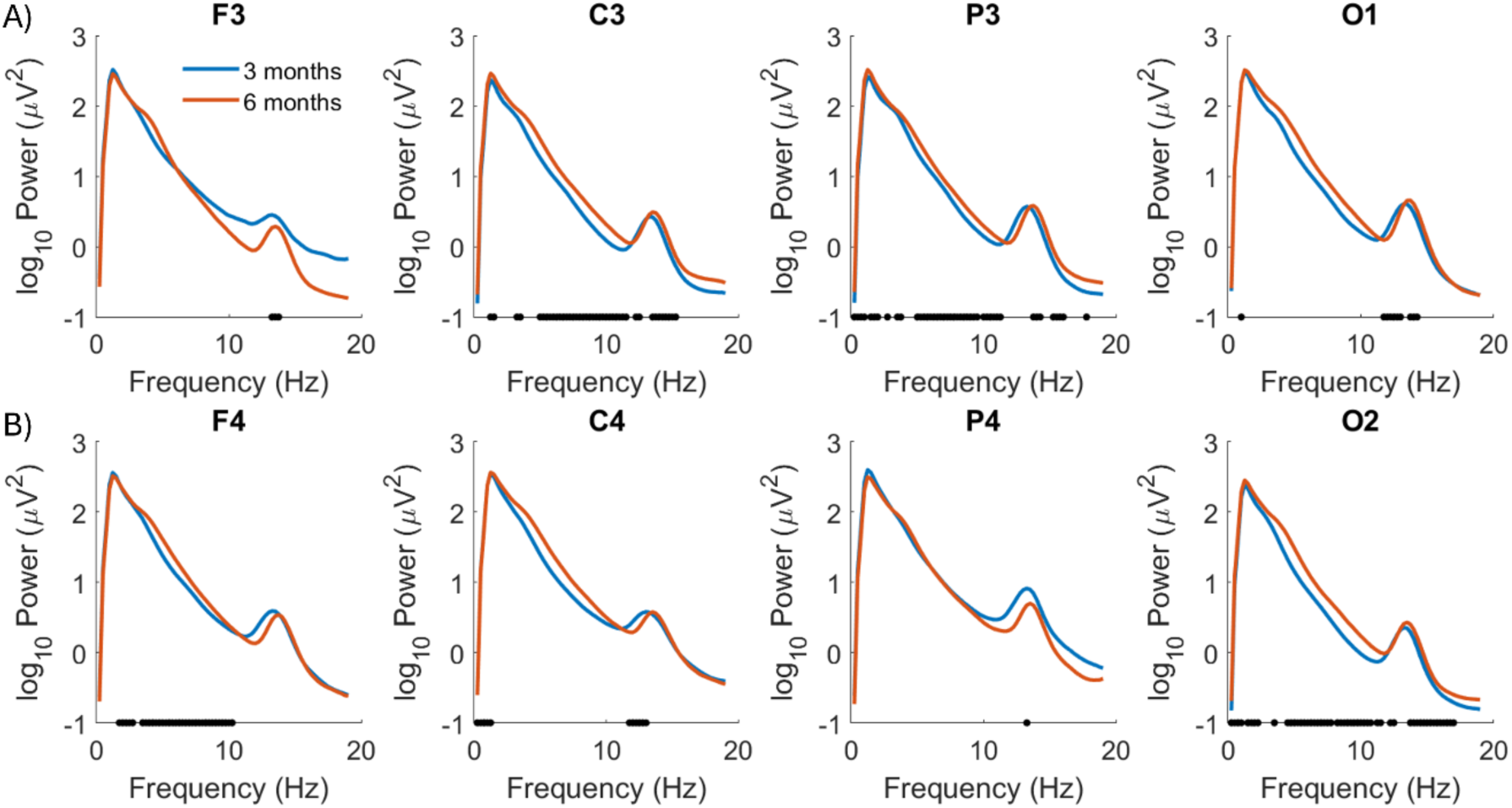
Power spectra averaged across infants for the 3-month (in blue) and 6-month (in red) assessments (n=9) including electrodes over A) the left hemisphere and B) the right hemisphere. The black dots indicate the frequency bins with significant differences between 3 and 6 months (corrected p<0.05).

In the right hemisphere, the frontal electrode (F4) showed a significant power increase from 3 to 6 months, mirroring the pattern observed in the left hemisphere (Figure 1B). However, hemispheric differences emerged centrally, where the right central electrode (C4) exhibited a significant increase in power across the 1-8 Hz frequency range. The right parietal electrode (P4) showed an even broader increase from 1-11 Hz, intensifying the maturation observed contralaterally. In contrast, the right occipital electrode (O2) showed power increases restricted to the 4-5 Hz and 14-19 Hz bands.

Overall, both hemispheres exhibited a maturational increase in NREM sleep EEG power, though with distinct regional patterns. The right hemisphere exhibited stronger central and parietal changes, while the left hemisphere showed greater occipital developmental dynamics.

Next, to explore individual differences in the spatial distribution of infant sleep EEG, we computed the topographical maps of SWA, theta power, and sigma power for all 11 subjects. The topographies revealed notable individual variability in the power distribution patterns across subjects (Supplementary Figure 1), with the coefficient of variation between subjects ranging from 18 to 80% (mean = 30% for SWA, 42% for theta power, and 46% for sigma power) at 3 months and from 16 to 67% (mean = 37% for SWA, 38% for theta power and 45% for sigma power) at 6 months. Prominent SWA was consistently observed over the occipital regions at both 3 and 6 months of age. In contrast, theta power exhibited a more diffuse distribution across the scalp, indicating emerging local patterns, particularly over the frontal and occipital regions. Sigma power demonstrated a strong central focus at 3 months that extended towards frontal regions by the age of 6 months. These insights highlight both the individuality and developmental trends in infant sleep EEG topography. Our quantitative approach confirmed these observations: we identified region-specific changes in topography maturation by analyzing the data averaged across the two age groups (Figure 2).

**Figure 2.**
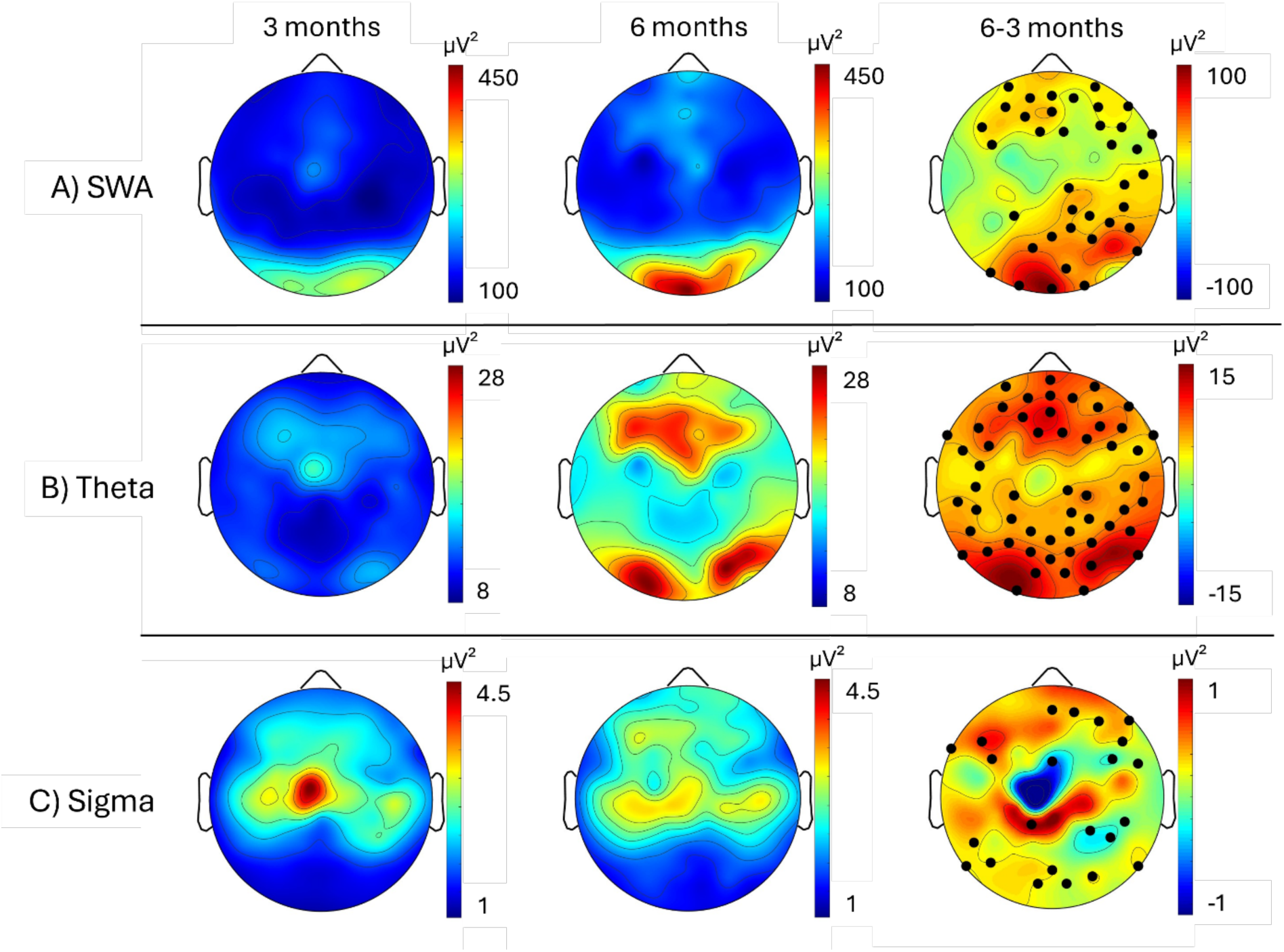
Topographical maps of (A) SWA, (B) theta power, and (C) sigma power averaged across infants for the 3-month (first column) and 6-month (second column) assessments (n=9). Age-related changes were calculated as the difference between power at 3 and 6 months (i.e., 3-month subtracted from 6-month data; third column). Electrodes showing significant age effects (corrected p < 0.05) are marked with black dots: 46 electrodes for SWA, 68 for theta power, and 35 for sigma power.

Beyond characterizing EEG topography in healthy infants, our findings highlight key developmental changes within this critical window. Electrode-wise comparisons revealed an increase in SWA from 3 to 6 months across 42% of electrodes by up to 65% (mean = 30%), with the most pronounced changes observed in occipital regions and extensions toward anterior areas (Figure 2A). Similarly, theta power showed a global increase across 62% of electrodes by up to 162% (mean = 80%), with the most prominent changes over frontal and occipital regions (Figure 2B). In contrast, sigma power exhibited a more localized shift across 32% of electrodes, with increases over frontal and posterior regions by up to 48% (mean = 26%) but confined central and parietal decreases by up to 13% (mean = 8%; Figure 2C). This suggests transitioning from a centralized sigma power distribution at 3 months to a more dispersed pattern at 6 months. Overall, our results indicate widespread maturational increases in sleep EEG power across frequencies, except for spatially confined decreases in sigma power.

Next, we analyzed the relationship between SWA, theta power, and sigma power maturation (i.e., power differences from 3 to 6 months) and behavioral outcomes at 6 months. For SWA power (Figure 3A), a positive correlation was observed over frontal regions with the Collective score (0.72 ≤ ρ ≤ 0.99, 0.007 ≤ p ≤ 0.048), Gross motor score (0.69 ≤ ρ ≤ 0.94, 0.004 ≤ p ≤ 0.049) and Personal social score (0.72 ≤ ρ ≤ 0.95, 0.01 ≤ p ≤ 0.043). This indicates that a greater increase in SWA from 3 to 6 months is associated with better performance across behavioral domains, albeit only locally. In contrast, a negative correlation with Personal social scores emerged in a very confined central area, indicating that centrally increased SWA is linked to less mature development in this domain (−0.95 ≤ ρ ≤ −0.8, 0.013 ≤ p ≤ 0.02).

**Figure 3.**
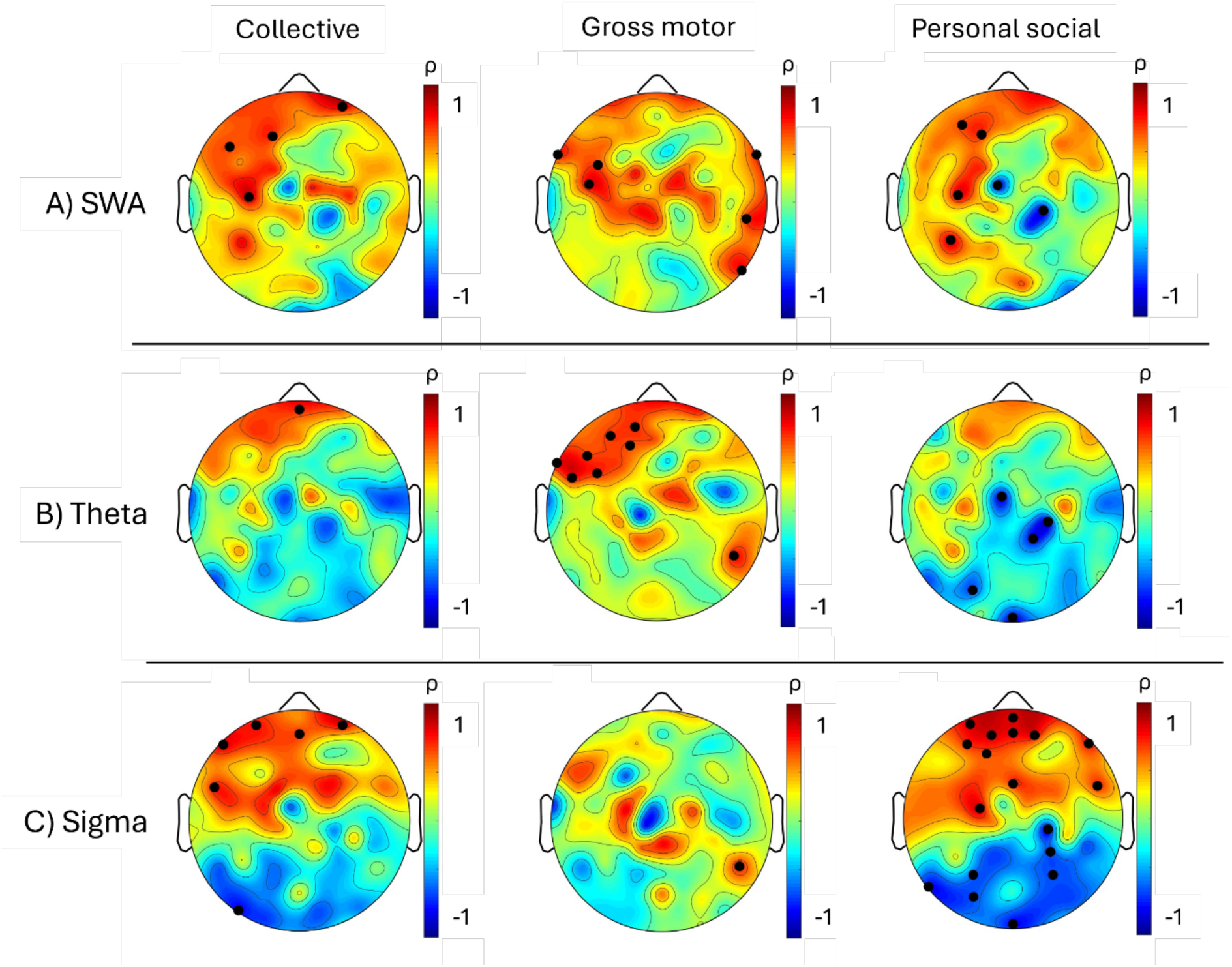
Topographical maps of Spearman correlations between changes in sleep EEG power from 3 to 6 months and behavioral scores at 6 months (Collective, Gross motor, Personal social). Correlations were computed for three frequency bands: A) SWA, B) theta, and C) sigma power. Significant electrodes are marked with black dots (corrected p < 0.05).

For theta power (Figure 3B), a positive correlation with the Collective score was identified at one frontal site (ρ = 0.81, p = 0.048). Gross motor scores were positively associated with theta power increases over left frontal areas (0.69 ≤ ρ ≤ 0.94, 0.002 ≤ p ≤ 0.043). In contrast, Personal social scores showed a negative correlation with theta power over posterior regions (−0.95 ≤ ρ ≤ −0.74, 0.01 ≤ p ≤ 0.036), indicating that a smaller increase in theta power in these regions was linked to better social skill development.

For sigma power (Figure 3C), positive associations were found between frontal sigma power and both the Collective (0.74 ≤ ρ ≤ 0.95, 0.015 ≤ p ≤ 0.048) and Personal social scores (0.7 ≤ ρ ≤ 0.96, 0.001 ≤ p ≤ 0.045). However, negative correlations were observed over occipital regions (ρ = −0.9, p = 0.019 for Collective; −0.95 ≤ ρ ≤ −0.68, 0.009 ≤ p ≤ 0.048 for Personal social). Additionally, Gross motor scores positively correlated with sigma power at one posterior site (ρ = 0.79, p = 0.024).

A consistent pattern emerged across frequencies and behavioral scores: increased frontal power was linked to more advanced skills. In contrast, posterior power increases correlated with lower scores, suggesting that a developmental shift in EEG power from posterior to anterior regions may indicate more advanced neurodevelopmental progress. To account for individual differences in baseline behavior, we additionally calculated partial correlations controlling for behavioral scores at 3 months. These analyses yielded similar topographical patterns (Supplementary Figure 2), suggesting that the observed associations are robust to individual differences in baseline behavior.

## 4. Discussion

This study provides the first longitudinal, high-density characterization of infant sleep EEG, enabling the detection of individual developmental trajectories across a sensitive window for plasticity. Our findings indicate that infant sleep neurophysiology evolves through both global and localized changes during this critical developmental period. Notably, SWA exhibited the most prominent increase in occipital regions, with power rising by up to 65%, and further frontal and parietal increases, likely reflecting early cortical restructuring on a broad scale. Meanwhile, theta power increased considerably across the entire scalp by up to 162%, reflecting broad neurodevelopmental progression. In contrast, sigma power, initially concentrated centrally, became more widely dispersed, suggesting a transition in thalamocortical connectivity. The identified region- and frequency-specific shifts delineate typical developmental trajectories and may reflect neuroplastic mechanisms guiding emerging skills, highlighting their potential as pediatric neurodevelopmental markers.

Our findings demonstrate a marked increase in SWA from 3 to 6 months, not only over occipital but also extending into frontal and parietal regions. While occipital dominance of SWA has previously been observed at 3 (Guyer et al., 2019) and 6 months (Schoch et al., 2021), our results fill the critical developmental gap between these time points, revealing the unfolding maturation of SWA topography during this period. The broader topographical expansion of SWA across infancy aligns with its known associations with white matter development in childhood, with SWA potentially reflecting early microstructural changes in myelin formation (LeBourgeois et al., 2019). The observed developmental SWA increase aligns with MRI-based trajectories of white matter expansion, following inferior-to-superior, anterior-to-posterior, and diagonal growth patterns from birth to 6 months (Grotheer et al., 2022; Kang et al., 2011). This pattern also coincides with the rapid synaptogenesis occurring in the occipital cortex during this period (Huttenlocher & Dabholkar, 1997; Natu et al., 2021). Given the role of SWA in synaptic homeostasis and plasticity (Tononi & Cirelli, 2006), our findings likely reflect active cortical remodeling during early infancy. Overall, these regional SWA increases not only align with cortical myelination but also support the concept that SWA may act as a mechanistic driver of early learning-related plasticity. These findings position infant SWA as a sensitive marker of neuronal network growth with potential for early diagnosis and intervention.

Our results show a global increase in theta power from 3 to 6 months, except for central and temporal regions. Theta activity during NREM sleep has been identified as a transitional marker of sleep homeostasis in infants, reflecting the build-up and dissipation of sleep pressure (Jenni et al., 2004).

Our findings refine this understanding by highlighting that theta power is not uniformly distributed across the cortex, in contrast to earlier studies reporting general age-related increases based on a limited number of EEG derivations (Jenni et al., 2004; Sankupellay et al., 2011). Beyond this, we here identify regional maturation that carries behavioral relevance, i.e. left frontal power was positively associated with gross motor function, and posterior power was linked to reduced personal-social skills, suggesting that regional theta maturation may reflect early differences in motor and social development. It remains exciting to be clarified whether theta power can further serve as a neurodevelopmental indicator concerning temperament, cognitive function, and behavioral regulation. Given that atypical theta activity has been linked to neurodevelopmental conditions such as ADHD (Castelnovo et al., 2022), future studies could assess its predictive value for identifying individual differences in early brain development and potential risk factors for later cognitive and behavioral outcomes.

We observed a shift in sigma power from a central to a more distributed pattern between 3 and 6 months, likely reflecting the maturation of spindle-related thalamocortical networks (Sanchez-Vives & McCormick, 2000; Steriade, 2006) and early plasticity processes linked to learning (Friedrich et al., 2019; Jaramillo et al., 2023). As sigma power matures, it mirrors changes in spindle dynamics, including frequency, density, and topographical spread shifts. Notably, sleep spindles appear by the age of 2 months, a developmental milestone accompanied by an increase in sigma power (Jenni et al., 2004). Between 3 and 6 months of age, spindle characteristics such as density and duration remain stable (Louis et al., 1992), whereas sigma power decreases, though this observation was limited to measurements from single EEG derivations over central and parietal areas (Ktonas et al., 1995; Sankupellay et al., 2011). Our findings build on this foundation and unveil remarkable regional variations in the maturation of sigma power, which are only discernible with high-density EEG approaches. These variations may be related to the region-dependent shift in GABAergic transmission from excitatory to inhibitory, a pivotal transformation that influences early neural activity and synapse formation in the first year of life (Murata & Colonnese, 2020; Tyzio et al., 2007). This shift has been proposed as a possible mechanism driving the maturation of sleep spindle features during this critical period (Chegodaev et al., 2022). Furthermore, thalamocortical connectivity within higher-order networks, particularly the default-mode and bilateral frontoparietal networks, has been shown to dramatically increase over the course of the first year (Alcauter et al., 2014). The observed evolution in sigma power, therefore, likely signals the progressive strengthening of thalamocortical circuits and a shift from short-range to long-range connectivity. This shift reflects the increasing integration of distributed cortical networks, a hallmark of early neural organisation and functional connectivity (Fair et al., 2007).

We found an association between the change in power from 3 to 6 months and behavioral development, such that greater power increases over frontal regions correlated with more advanced skills at 6 months across frequencies and behavioral domains. This was particularly pronounced for gross motor skills in the theta band and personal social skills in the sigma band. Interestingly, smaller increases or even reductions in power over posterior regions from 3 to 6 months corresponded with more advanced skills at 6 months, a pattern especially noticeable for personal social skills in the sigma band. Given that synaptogenesis in the prefrontal cortex begins in the fetus but progresses more gradually than in the visual cortex and peaks after the first year, in mid-childhood (Huttenlocher & Dabholkar, 1997; Liu et al., 2012), our findings underscore the link between anteriorization (Novelli et al., 2016) and skill maturation. This relationship reveals a subtle yet functionally significant maturation in the frontal regions during this early developmental period, which often remains obscured due to the more pronounced changes occurring in posterior regions. Our findings suggest that longitudinal changes in sleep EEG power may be a more significant marker of skill maturation than previously examined concurrent power levels (Guyer et al., 2019; Page et al., 2018; Satomaa et al., 2020). This approach uniquely captures time-sensitive neurophysiological shifts that precede behavioral expression, revealing the early unfolding of plasticity in the developing brain. The previously reported predictive nature in the relationship between sleep spindle activity at 6 months and behavioral outcomes at 12 and 24 months (Jaramillo et al., 2023) supports the notion that sleep EEG activity reflects brain maturation processes essential for later behavioral development. This provides valuable insights into how early neural dynamics may predict later developmental outcomes.

Despite their potential, sleep-related markers of early neurodevelopment remain underutilized. Many neurodevelopmental disorders are not identified until school age (Bachmann et al., 2017; Brett et al., 2016; Sheldrick et al., 2017), underscoring the need for early biomarkers to detect atypical development before symptoms emerge. While most research relies on single time-point assessments, longitudinal sleep EEG data offer more reliable maturation markers by capturing individual variability and enhancing statistical power (Bruni et al., 2022). Tracking neurophysiological transitions over time is essential for a comprehensive understanding of neural network maturation, helping to identify early indicators of developmental risks. Therefore, we call on future research to refine sleep-related EEG biomarkers for diagnostics and intervention by prioritizing two key aspects: longitudinal data and clinical populations. Only with this focus can we enable early detection and personalized interventions that may meaningfully alter the trajectory of neurodevelopmental disorders, ensuring timely and effective support for those affected.

When interpreting our findings, the small sample size must be considered as a limitation. A limited dataset can inflate correlation coefficients, potentially overestimating the strength of associations, particularly when using traditional correlation methods. To address this, we conducted a secondary analysis using biweight midcorrelation, a technique that is less sensitive to outliers and provides a more robust measure of association. This analysis confirmed our results’ consistency, strengthening our conclusions’ reliability. Studies of infant sleep are often constrained by sparse electrode sampling and small numbers of subjects (Chu et al., 2014). Yet, even in small samples, longitudinal sleep EEG measures have shown predictive value for later developmental outcomes (Kurth et al., 2013; LeBourgeois et al., 2019). Nevertheless, future studies with larger populations are needed to validate these findings and ensure their generalizability.

## 5. Conclusion

This study delineates the topographical maturation of infant sleep EEG from 3 to 6 months, revealing both local and global neurophysiological changes. We identified a pronounced increase in occipital, frontal and parietal SWA, a widespread rise in theta power, and a dispersion of sigma power from originally central locations. Crucially, these changes correlated with motor and social skill development, highlighting the prognostic value of sleep neurophysiology for emerging behavior and brain function. This underscores the infant sleep EEG as a non-invasive biomarker to predict and monitor individual developmental trajectories even before overt behavioral signs appear. By capturing early cortical dynamics linked to brain plasticity, infant sleep EEG can inform diagnostics, guide personalized interventions, and support the evaluation of treatment in preverbal populations. This work lays a foundation for clinical applications of sleep neurophysiology in early identification of atypical development and advances our understanding of neuroplasticity during infancy.

## Acknowledgments

We are especially grateful to the families who participated in the study for their time, trust, and patience. Their commitment made this research possible. We thank Estelle Bader and Leila Bajrami for their help with data collection and Sara Dozio for her support with graphics.

---

**Supplementary Figure 1.**
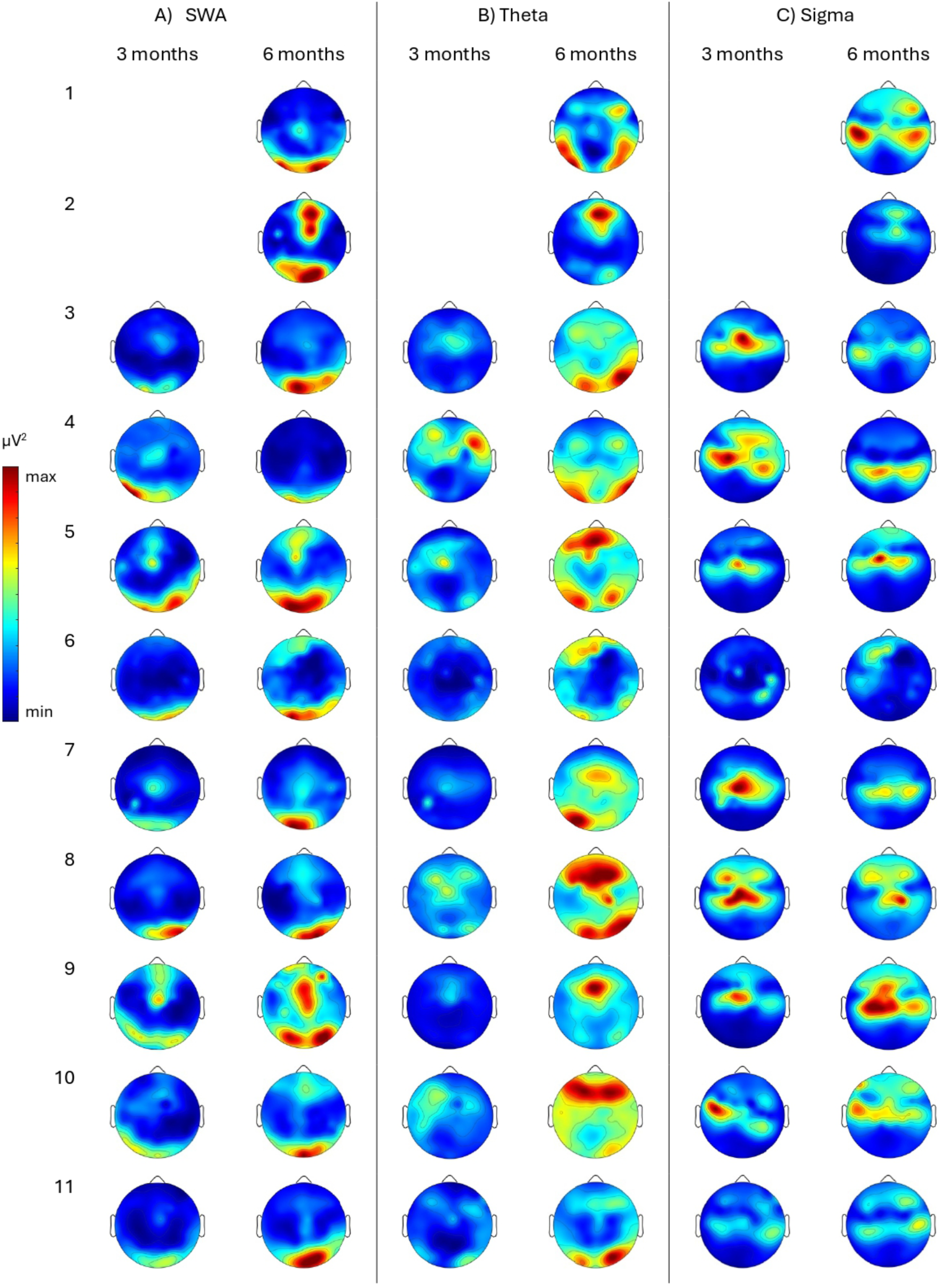
Individual topographical maps of (A) SWA, (B) theta power and (C) sigma power for each infant at the 3-month (first column) and the 6-month (second column) assessment (n=11 with 6-month assessment and n=9 with both assessments). Each row represents a subject, with missing topographies indicating missing data. The maps belonging to the same infant are plotted on a consistent scale within a frequency band.

**Supplementary Figure 2.**
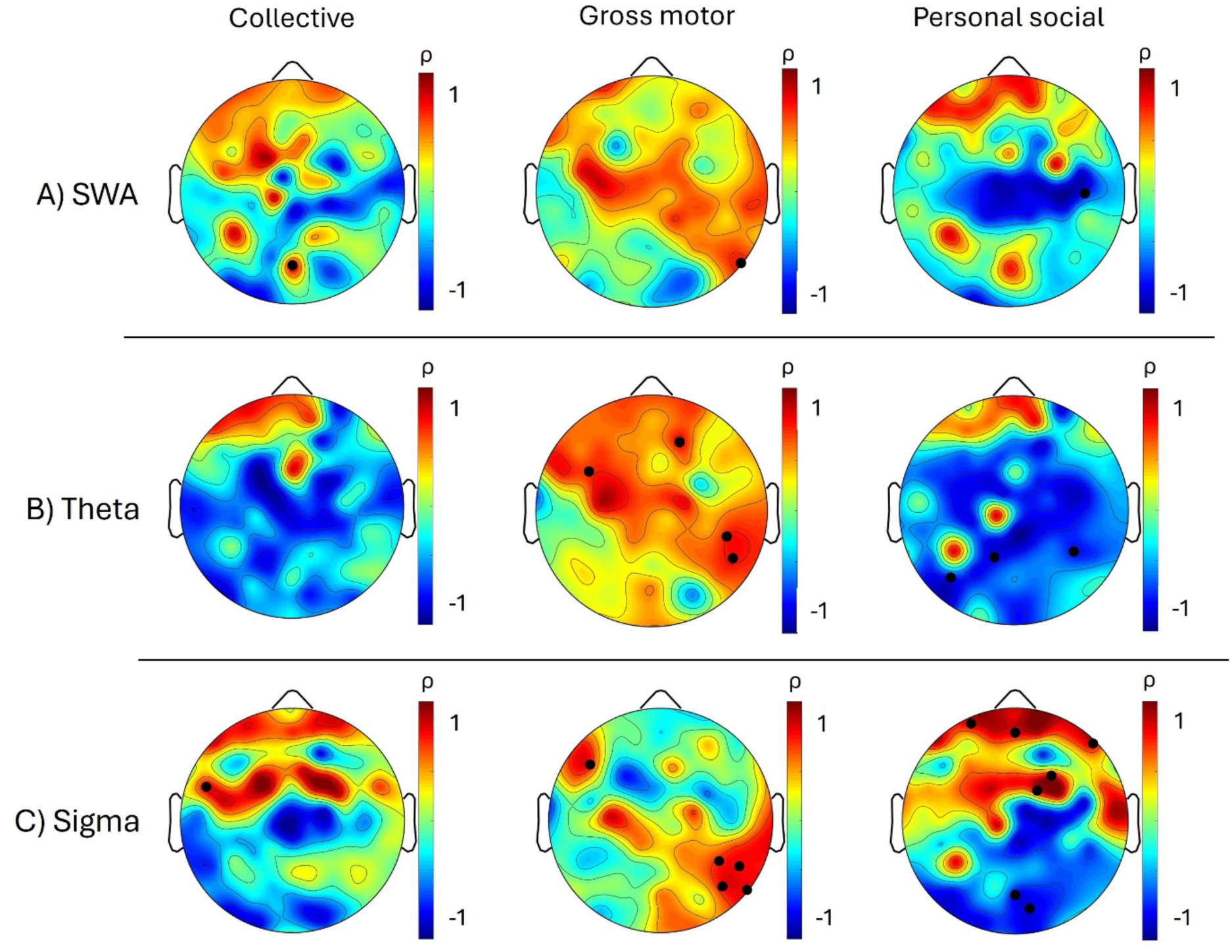
Topographical maps of partial Spearman correlations between changes in sleep EEG power from 3 to 6 months and behavioral scores at 6 months (Collective, Gross motor, Personal social), controlling for behavioral scores at 3 months. Correlations were computed for three frequency bands: A) SWA, B) theta, and C) sigma power. Significant electrodes are marked with black dots (corrected p < 0.05).

